## Additional file2: Supplement figures for "MalariaSED: a deep learning framework to decipher the regulatory contributions of noncoding variants in malaria parasites"

Fig. S1

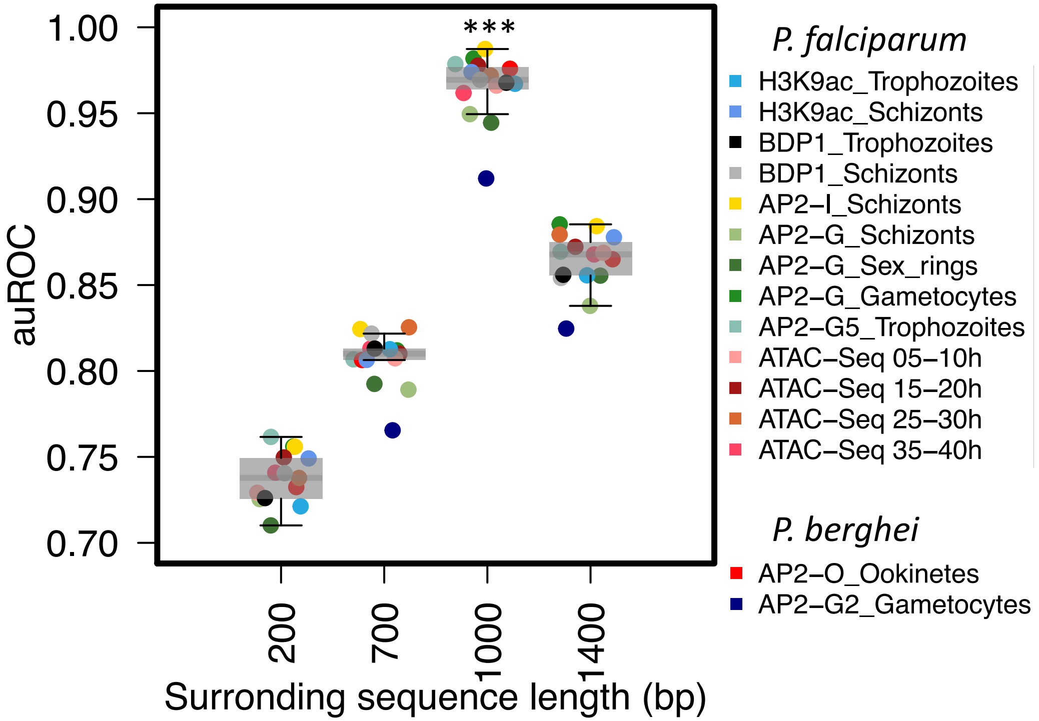


**Fig. S1. Evaluating the performance of MalariaSED using different lengths of DNA sequence input.** The same DL architecture as MalariaSED was trained on 200, 700, 1000 and 1400 bp DNA sequences, respectively. We present the auROC of each model as a point and use a box plot to show the distribution of the models inputting the same DNA sequence length. The results indicate 1000bp input outperforms other length input ('***' indicates Wilcoxon test $p < 0.01$ compared between the 1kb input sequence and any other length listed here ).

Fig. S2

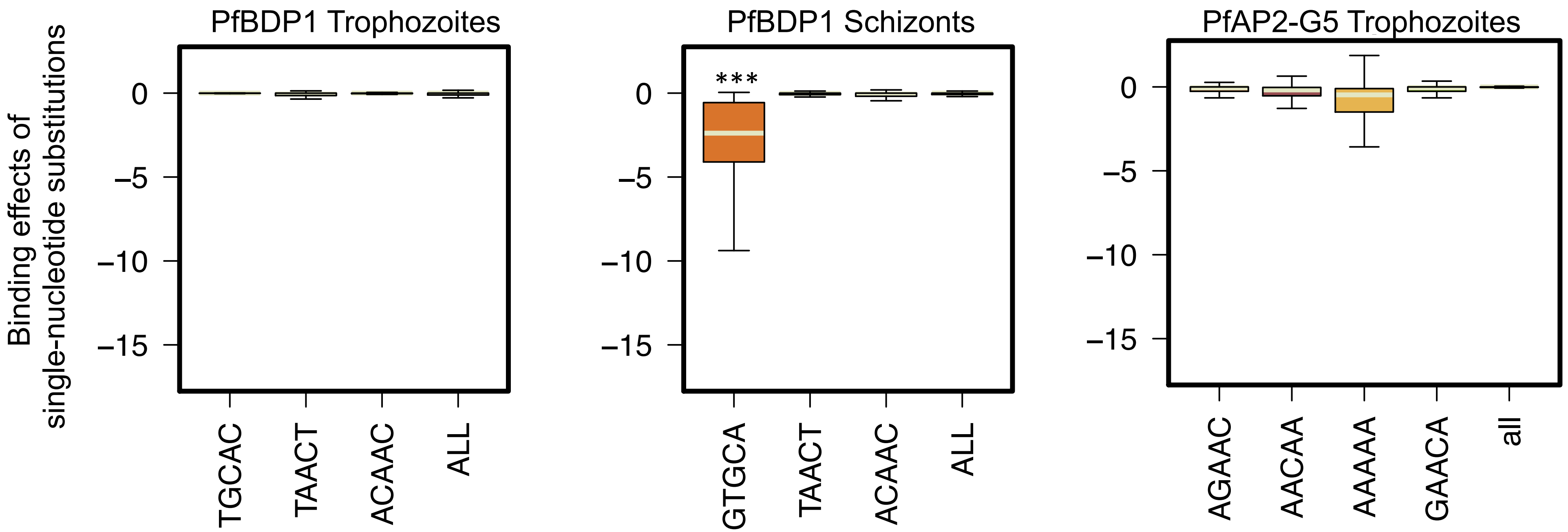


**Fig. S2. TFs binding effects of single nucleotide substitutions at 5-mers sequences predicted by MalariaSED in *P. falciparum.*** '***' indicates Wilcoxon test compared with whole-genome background $p < 2.2e-16$.

Fig. S3

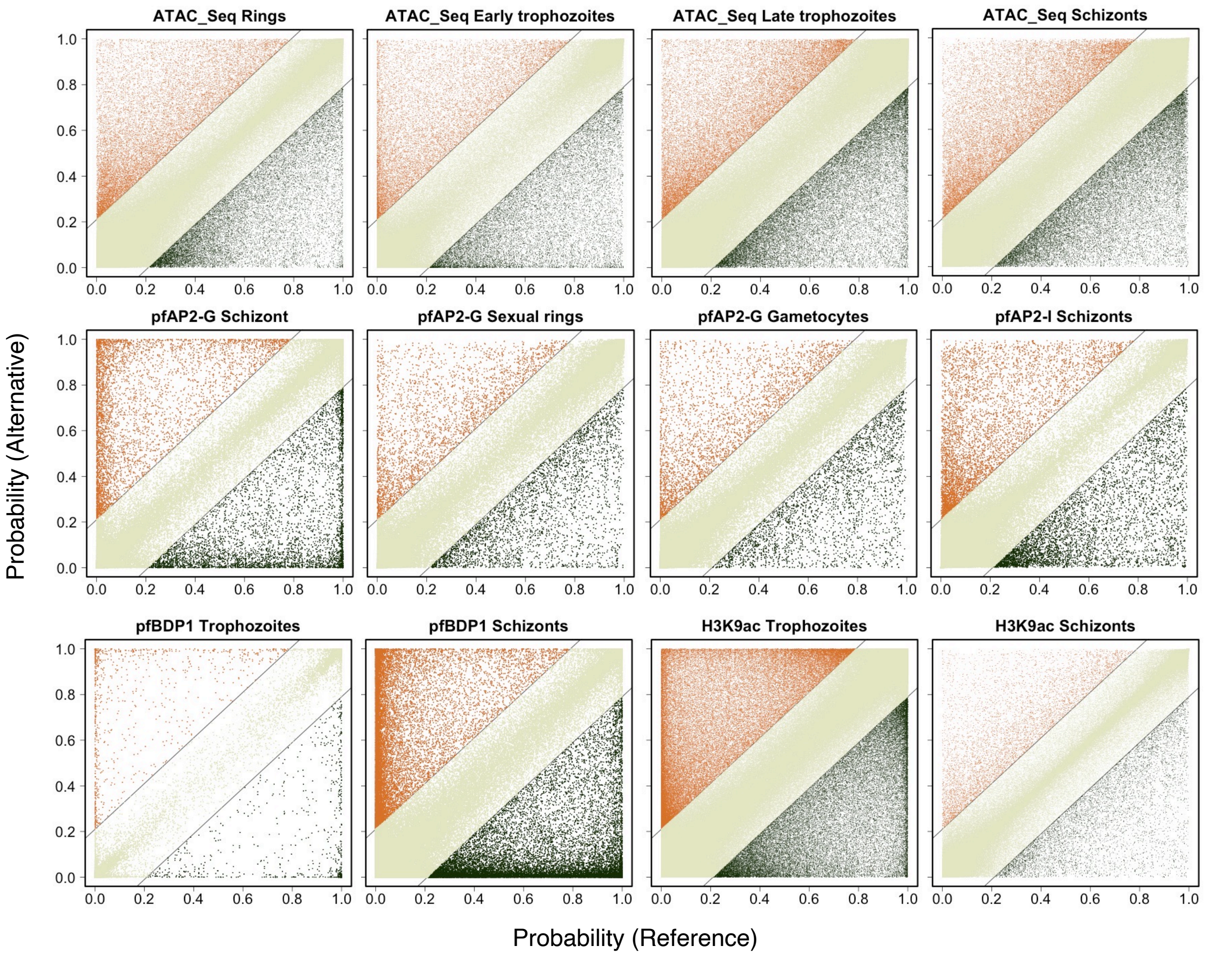


**Fig. S3. MalariaSED prediction for epigenetic markers of ~1.3 millions variants gathered by Pf3K.** The horizontal and vertical direction represent, respectively, the predicted probabilities that the sequences carrying the reference allele and alternative allele.

Fig. S4

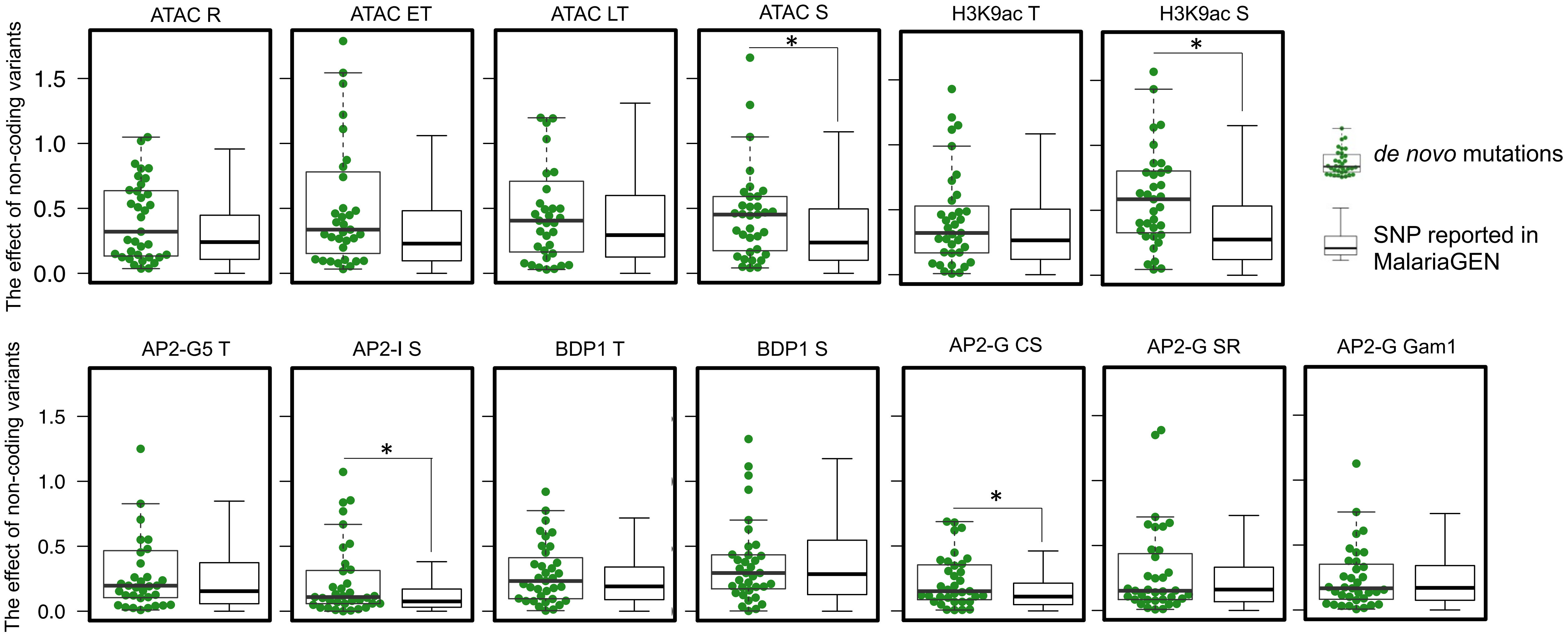


**Fig. S4. The results from MalariaSED indicates de novo mutations at non-coding regions discovered from the single cell study have higher chance to alter their surrounding chromatin profiles in schizonts.** '*' indicates Wilcoxon test compared with non-coding SNPs reported from Pf3K, $p < 0.05$.

Fig. S5

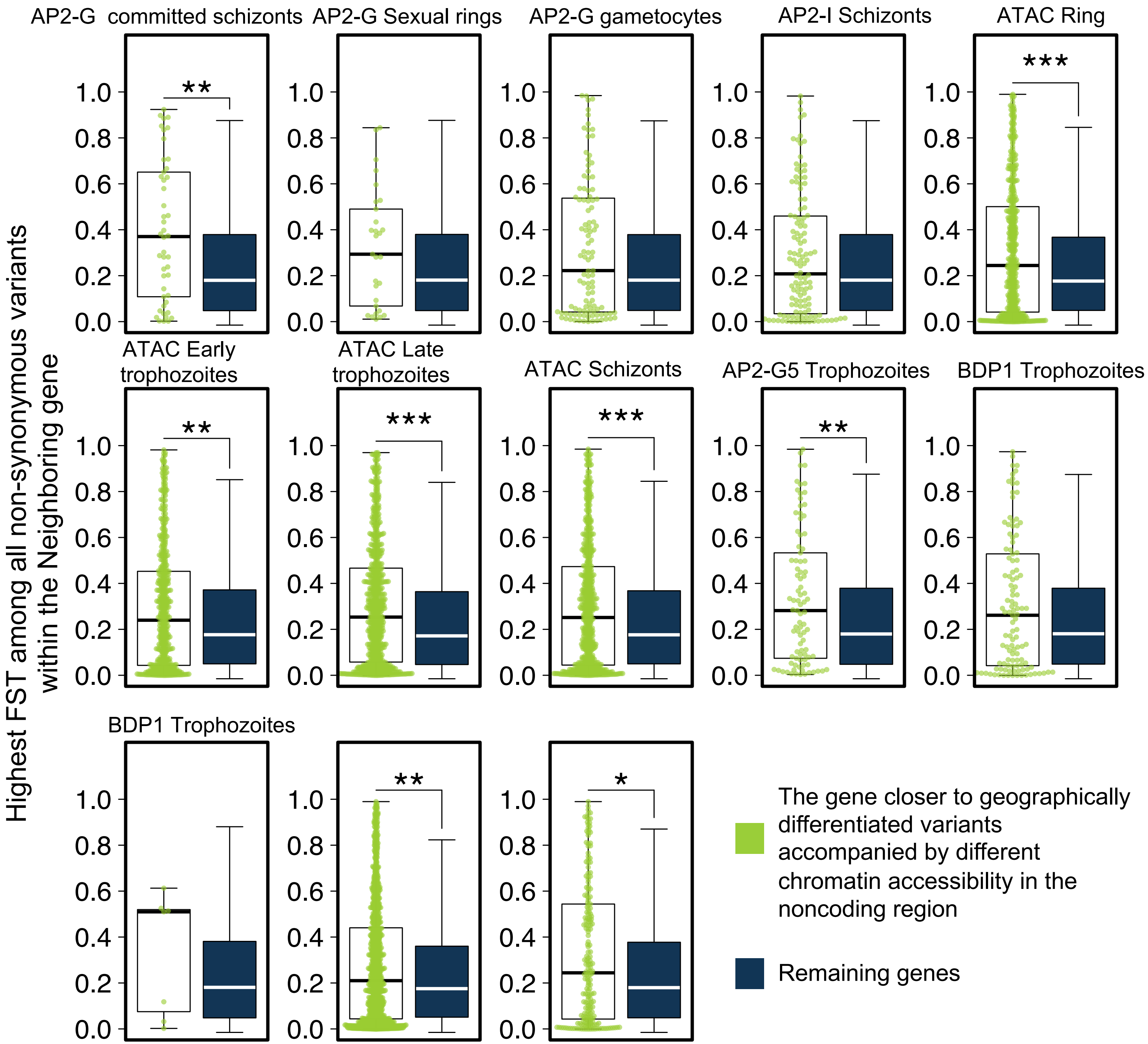


**Fig. S5. The geographically differentiated variants accompanied by different chromatin profile effects in the noncoding region are more likely closer to genes with high levels of geographic differentiation.** Each green dot represents a gene closest to noncoding variants presenting global FST > 0.1 and high chromatin effects (the top 1% of chromatin effects). '*' represents Wilcoxon test p< 0.05, '**' is p < 0.01, while '***' < 1e-3.
